# A synergistic culture dependent and independent approach reveals a conserved wheat seed mycobiome

**DOI:** 10.1101/2024.02.22.581674

**Authors:** Lindsey E. Becker, David Marshall, Marc A. Cubeta

**Affiliations:** Department of Entomology and Plant Pathology, Center for Integrated Fungal Research, North Carolina State University, Raleigh, NC 27695, USA; USDA-ARS Plant Science Research Unit, Department of Entomology and Plant Pathology, North Carolina State University, Raleigh, NC 27695, USA

## Abstract

The occurrence of pathogenic fungal taxa associated with wheat (*Triticum aestivum* L.) seeds is well studied, but less is known about non-pathogenic taxa of the wheat seed mycobiome. The goal of our research is to characterize wheat seed fungal endophyte diversity with a synergistic culture dependent and independent experimental approach. Four publicly available winter wheat cultivars developed in the southeastern United States with varying phenotypic and disease resistance traits were examined over a period of two years: Catawba, Hilliard, Shirley, and USG 3640. Our culture dependent methods involving two nutrient media generated 645 fungal isolates representing twelve genera sampled from multiple cultivars. Metabarcoding analysis identified a broader range of fungal taxa and a greater number of unique sequences than culture dependent methods. When examining fungal diversity across cultivars and years, richness decreased in 2021 for both culture dependent and independent approaches. However, wheat seed fungal community structure was stable across cultivars and years. Our results highlight the importance of combining culture independent and dependent methods to capture and establish a diverse endophytic fungal catalog associated with the wheat seed and highlight areas where future culture dependent efforts can focus their efforts.

## INTRODUCTION

Bread wheat (*Triticum* spp.) is a highly adaptable staple crop grown globally and provides nearly a fifth of the world’s available calories and protein (Shiferaw *et al*., 2013). The advent of the Green Revolution led to enormous improvements in yield for wheat farmers, however over the past decade improvements in grain yield have staggered (Calderini *et al*., 2021). With continuously growing global demand, slight variation in wheat production can lead to increases in overall food prices. The United States in 2021 experienced a 10% decline in wheat yield compared to the year before even with planted acreage increasing by 1% (USDA-NASS, 2021). This decrease was largely attributed to droughts and heatwaves (Sowell & Swearingen, 2021). Climate variability and its negative impacts are expected to intensify in coming decades (Zhao *et al*., 2017). There is a need to identify novel strategies to aid plant adaptation to a rapidly changing environment. Plant associated microbial communities, or microbiomes, represent a sustainable tool for crop resilience and exploiting this resource with other management tools will support crop production. Plant pathogens associated with crop seeds have been implicated in reduced storage life, crop failure, and phytosanitary restrictions (Singh & Mathur, 2004; Munkvold, 2009; Choudhury *et al*., 2017). Seedborne pathogens receive much attention and are well monitored in crop production (Gitaitis & Walcott, 2007). However, fungi that reside within the seed without causing visible harm have received little attention despite the increased desire to understand the influence of the microbiome on plant host success (Nelson, 2018; Gdanetz *et al*., 2021). Endophytic fungi, harbored inside plant seeds, evolved to establish intimate and long lasting relationships with the plant host that can yield negative, neutral, or positive effects (Schardl *et al*., 2004; Schulz & Boyle, 2005; Matanguihan *et al*., 2011). Both unicellular and multicellular fungi play important roles in the seed microbiome (Vujanovic *et al*., 2012; Mascot-Gómez *et al*., 2021; Bintarti *et al*., 2022).

Fungal communities occupy seeds through waves of introductions from other plant organs and surrounding environments (Larran *et al*., 2007; Comby *et al*., 2016; Hertz *et al*., 2016). Assembly of the seed microbiome is generally understood as occurring through vertical (maternal) and horizontal (environmental) transmission (Hardoim *et al*., 2015; Shade *et al*., 2017). Maternal transmission can result from direct or indirect entry through vascular and nonvascular tissue from the mother plant into progeny seed. Common pathogenic fungal taxa such as *Fusarium* and *Verticillium* represent classical examples of maternal transmission via vascular tissue (Singh & Mathur, 2004). Gagic *et al* (2018) highlighted the variability of seed-borne transmission of *Epichloë* species in both seeds and primordia of ryegrass with successful transmission influenced by host genotype. Indirect entry through floral tissues has also been well documented in the cereal ergot disease caused by *Claviceps purpurea* (Singh & Mathur, 2004). Horizontally transmitted fungi often localize and aggregate mycelial growth on the seed surface and seed coat (Pochon *et al*., 2012). Histopathological experiments revealed that *Cercospora sojina* infects soybean seed surface and penetrates the seed coat through pores and cracks (Singh & Sinclair, J.B., 1985; Agarwal & Sinclair, 2014). In terms of microbial transmissibility in plant seeds, microbiome studies lack the ability to examine vertical or horizontal passing as strain level differentiation of ASVs remains limited (Callahan *et al*., 2021). Instead, these studies rely on co-occurrence of ASVs to predict transmission pathways for taxa of interest (Bergna *et al*., 2018; Abdelfattah *et al*., 2021; Latz *et al*., 2021). Overall, there is much that remains to be understood regarding the assembly, transmission and function of the seed microbiome in relation to plant host success.

The bulk of wheat microbiome studies focus on microbial communities associated with roots and leaves, while very few aim to define the seed microbiome (Karlsson *et al*., 2017; Gdanetz & Trail, 2017; Kavamura *et al*., 2018; Mavrodi *et al*., 2018; Schlatter *et al*., 2020; Hassani *et al*., 2020; Simonin *et al*., 2020). Wheat seed microbiome studies generally prioritize examination of bacterial communities over fungal, viral and protozoal communities (Robinson *et al*., 2016; Díaz Herrera *et al*., 2016; Kavamura *et al*., 2019; Kuźniar *et al*., 2020; Hone *et al*., 2020; Walsh *et al*., 2021). The earliest examination of the wheat seed fungal endophytes relied on culture-dependent methods, often with an emphasis on isolating pathogenic taxa and recovering a limited number of isolates (Christensen, 1957; Flannigan, 1974; Sieber *et al*., 1988; Larran *et al*., 2007). The most commonly isolated fungal endophytic taxa included *Alternaria*, *Epicoccum*, *Fusarium*, *Stagonospora*, and *Cladosporium* (Sieber *et al*., 1988; Comby *et al*., 2016). While culturing methods are known to capture a small subset of plant associated fungi, fungal isolate collections provide invaluable resources for researchers studying the plant microbiome. Fungal isolates provide opportunities to improve our understanding of plant-associated fungi both *in vitro* and *in planta* (Anguita-Maeso *et al*., 2020). Isolate collections can allow for better genomic resolution via multi-gene or whole genome sequencing (Dissanayake *et al*., 2018). With whole genome sequences of seed associated fungal isolates, researchers can develop strain specific primers to examine transmission, monitor abundance and record persistence over generations (Wassermann *et al*., 2021). With the advent of next generation sequencing, the number and phylogenetic range of unculturable fungal taxa associated with wheat seeds has dramatically increased (Bakker & McCormick, 2019; Rojas *et al*., 2020). Comparison of seed microbiomes across cultivars, locations, resistance to disease, and generations has slowly begun to provide snapshots of how these factors influence the seed microbiome (Larran *et al*., 2007; Bakker & McCormick, 2019; Rojas *et al*., 2020; Latz *et al*., 2021). However, both culture dependent and culture independent analysis across genotypes and over multiple generations in the field could provide a more comprehensive understanding of the wheat seed mycobiome to a higher taxonomic resolution.

Given their importance as founding members of the plant microbiome, and the complex niche they inhabit in comparison to other plant organs, seed endophytic fungal communities require diverse approaches that combine both culture dependent and independent methods. Endophytic fungi have the potential to buffer the effects of a rapidly changing climate where too much rain and mild droughts are both predicted to escalate. The goal of this study is to elucidate the influence of wheat cultivars over two field seasons on the seed mycobiome. Specifically, we aimed to culture seed endophytes from four different winter wheat cultivars bred for the southeast United States; Catawba, Hilliard, Shirley, USG-3640, over a period of two years (2020 and 2021) and complement culture-based with culture-independent investigation of wheat seeds. The culture dependent efforts allowed us to build a reference library of cultures that were identified to the species level. Our culture independent investigation enabled us to determine alpha and beta diversity of fungal communities in wheat seeds and identify a core wheat seed mycobiome.

## MATERIALS AND METHODS

### Wheat Cultivars

Four winter wheat (*Triticum aestivum* L.) cultivars, Catawba, Hilliard, Shirley, and USG-3640, all bred and commonly grown throughout the southeastern United States were used in this study. Three of the cultivars, Hilliard, Shirley, and USG 3640, are classified as soft red winter wheat, while Catawba is classified as hard red winter wheat. Awn types varied by cultivar, with Hilliard and USG 3640 exhibiting awns, while Shirley is awnletted and Catawba is awnless. Cultivars also varied by heading date, with Catawba heading early, USG 3640 heading mid-season, with Hilliard and Shirley heading late season. Cultivars were largely resistant or moderately resistant to powdery mildew and leaf rust diseases, but varied in susceptibility to other common diseases such as head scab and glume blotch (Griffey *et al*., 2010, 2020; Mergoum *et al*., 2022).

### Field Experiments

Untreated seed of each cultivar sourced from Kinston, NC (Catawba, Hilliard, and Shirley) and Plains, GA (USG-3640) in summer of 2019 was planted in the fall of 2019 and 2020 at the MidPines Research Station, Raleigh, NC, USA. Fertilization followed standard recommendations for winter wheat cultivation in North Carolina and no fungicides were applied to plants in the field. Soil type at MidPines Research Station is classified as Cecil clay loam. Five replicate experimental plots (15.2 m by 1.5 m) were planted for each cultivar at a seeding rate of 3 seeds/sq m. A total of 20 plots were seeded in four rows in a completely randomized block design. Mature seeds were harvested from each plot in June with a combine. Seed harvested from each experimental plot was cleaned and weighed. Seed moisture content, protein, and test weight (kg/bushel) were determined.

### Seed Processing

Ten grams of seed of each cultivar harvested in 2020 and 2021 were disinfested by placing them in a sterile stainless steel tea strainer and dipped sequentially in 95% EtOH for 60 s, 3% active NaClO with 20 µL of Tween 20 (Thermo Fisher, Waltham MA, US) for 120 s, 95% EtOH for 30 s, sterile distilled H_2_O for 30 s, and sterile distilled H_2_O for 5 s. Seeds were air-dried inside a laminar flow hood. For culturing, 5 g of seed from each experimental plot were macerated in a sterile mortar and pestle in a laminar flow hood. The resulting seed fragments were sieved using Bel-Art Mini-Sieve (Bel-Art, Wayne NJ, USA), with a size 80 sieve that removed fragments greater than 177 µm. Sieved fragments were diluted 1:100 with sterile distilled H_2_O and gently vortexed before adding 500 µL of the wheat fragment solution to 9-cm diam nutrient media plates and dispersed across the plate using a plate spinner and sterile plate spreader. Three replicate plates were prepared for each cultivar and nutrient medium.

### Media Selection

Two nutrient media were selected for fungal isolation; Dichloran Glycerol (DG18) Agar and Malt Yeast Extract Agar (MYEA). DG18 is useful for isolating fungi that grow at low water potential and was prepared by adding 31.5 g of DG18 Agar (Oxoid, Waltham MA, US) with 220 mL of glycerol, 0.01 g ZnSO_4_, 0.005 g CuSO_4_, 500 µL of streptomycin (Fisher Scientific, Waltham, MA) at 100 mg/mL, and 500 µL of ampicillin (Fisher Scientific, Waltham, M) at 100 mg/mL, per L of distilled H_2_O. MYEA is a generalist selection medium for fungi and was prepared by adding 10 g of Malt Extract broth (BD Difco, Franklin Lakes, NJ) with 2 g of Yeast Extract (Fisher Scientific, Waltham, MA), 15 g of Agar (Fisher Scientific, Waltham, MA), 500 µl of streptomycin at 100 mg/mL, and 500 µL of ampicillin at 100 mg/mL. A second medium of MYEA was amended with 4 µL of cyclosporine at 1g/100 mL, a fungistatic compound that reduces hyphal expansion to isolate slower growing fungi that may be outcompeted by copiotrophic fungi (Bills *et al*., 2004; Pirttilä & S, 2011).

### Pure Culture Isolation and Morphotype Selection

Isolation plates were monitored for fungal colony growth every day for two weeks. The number and colony morphotypes were observed and recorded. Unique colonies were aseptically transferred from isolation plates to new MYEA plates. Cultivar and plate source were recorded at time of transfer, allowing for new isolates to be traced back to source. Once transfers were complete, isolates were grouped by macroscopic morphological characteristics. One representative from each morphological group was selected for DNA extraction.

### DNA extraction from pure cultures and Sanger sequencing

Pure cultures of each morphotype were transferred to a Malt Yeast extract broth and incubated for three weeks at room temperature. At three weeks, liquid broth cultures were filtered using WhatMan #10 filter paper (Cytiva, Marlborough, MA), vacuum pumped to remove excess liquid, with the remaining fungal tissue placed in 2 mL microcentrifuge tubes with two sterile 3-mm glass beads and lyophilized for 72 h with a Labconco benchtop freeze dryer (Labconco, Kansas City, MO). Lyophilized tissue was homogenized with an Omni bead mill (Omni, Kennesaw, GA). Tubes were shaken at a rate of 5 m sec^-1^ for 30 sec. Nucleospin Plant II mini kit (Macherey Nagel, Duren Germany) was utilized to extract DNA from pure cultures, adding the optional chloroform step for optimization of fungal DNA extraction. DNA concentration of pure cultures was measured using Qubit Fluorometer (Thermofisher, Waltham MA, US). ITS1 primers ITS1-F_KYO2 (TAGAGGAAGTAAAAGTCGTAA) and ITS86R (TTCAAAGATTCGATGATTCAC) were selected based on minimal amplification of *Triticum* genomic DNA from leaves and comparable amplification of fungi associated with the Ascomycota and Basidiomycota phyla, respectively (Scibetta *et al*., 2018). PCR amplification of the ITS1 region was conducted in a T100 Thermocycler (Bio-Rad, Hercules, CA US) with the following protocol for 35 cycles: 30 s at 98 C, 30 s at 56 C, 1 min at 72 C, and 10 min at 72 C. The master mix for the PCR reaction included 10 µL GoTaq Hot Start Green Master Mix (Promega, Madison, WI US), 2 µL of each primer at 10 µM, 6 µL of sterile water, and 2 µL of DNA template for a total of 20 µL reaction volume. Sanger sequencing was employed to sequence the ITS1 region of culture morphotypes. The CLC Genomics Workbench was used to visually inspect sequences and remove ambiguous bases. Sequences were placed by using the internal transcribed spacer region (ITS) from Fungi type and reference material using BLASTn option at https://blast.ncbi.nlm.nih.gov/ with attribution of genus and species provided when 97% or greater sequence similarities were available.

### DNA extraction and PCR amplicon sequencing from wheat seeds

For DNA extractions from seed, one surface disinfested seed was placed in a bead beating tube Type G (Macherey Nagel, Duren Germany) and shaken at a rate of 5 m sec^-1^ for 30 sec in a Bead Ruptor (Omni, Kennesaw, GA). Four seeds per cultivar per year were analyzed. Remaining disinfested seeds were stored at -20 °C in sterile microcentrifuge tubes. DNA extraction from the seeds and quantification was the same as described above for our culture approach. ITS1 primers described in culture dependent approach were used for amplicon sequencing, with slight modifications made for our PCR protocol for amplicon sequencing. Briefly, we executed a touchdown PCR in a T100 thermocycler, with the reaction volumes and cycling parameters detailed below (Bio-Rad, Hercules, CA). Total reaction volume was 50 µl, containing 2 µl of genomic wheat seed DNA at 50 ± 25 ng/µl, 25 µl DreamTaq PCR Master Mix 2X (Thermo Fisher Scientific, Waltham, MA), 2.5 µl of each primer (10 µM), and 18 µl of sterile H_2_O. PCR cycling conditions included two phases, an initial touchdown phase followed by an amplification phase. The touchdown phase involved denaturation at 95°C for 3 min; followed by 15 cycles of denaturation at 95℃ for 30 s, annealing at 63℃ for 45 s, and extension at 72℃ for 1 min, with annealing temperature decreasing by 1℃ for all 15 touchdown cycles. The generic amplification stage consists of 20 cycles of denaturation at 95℃ for 30 s, annealing at 56℃ for 45 s, extension at 72℃ for 1 min, with a final extension at 72℃ for 5 min (Korbie & Mattick, 2008). Amplification of PCR products was confirmed with 2% electrophoresis gels and visualized with ethidium bromide staining. PCR amplicon size was determined with a 100 bp DNA ladder. Library preparation and DNA sequencing were conducted at Michigan State Genomics Core. Sequencing was performed for 32 total samples in MiSeq v2 250 bp paired-end format (Illumina, San Diego, CA).

### Amplicon Sequence processing

To process raw reads, we employed DADA2 ITS Pipeline Workflow 1.8 parameters as standard, except for the following filtering parameters: maxN=0, maxEE=c(2,2), truncQ=2, minLen=50, rm.phix=TRUE and truncLen=c(190,150). Further steps such as denoising, merging, and filtering of chimeric reads were performed as standard within the workflow (Callahan *et al*., 2016). To prevent over inflation of diversity estimates due to utilization of Amplicon Sequence Variants (ASVs), we performed Lulu curation (Frøslev *et al*., 2017). Taxonomy assignment of curated ASVs with the UNITE ITS database Version 9.0 was performed in DADA2 (Abarenkov *et al*., 2010). All samples passed quality filtering parameters and were kept for downstream analysis.

### Statistical Analysis

Statistical analysis was conducted using the R environment (version 4.2.2) (R Core Team, 2018). Relative abundance graphs were produced in phylosmith (version 1.0.6) (Smith, 2019). Alpha diversity estimates for species richness were calculated in phyloseq (version 1.42.0), with analysis of variance (ANOVA) performed in vegan (version 2.6-4), with results plotted using ggplot2 (version 3.4.0) (Oksanen *et al*., 2022). Metacoder (version 0.3.5.00) was utilized to construct a phylogenetic tree of taxa captured within our culture independent approach (Foster *et al*., 2017). We utilized the package MicEco (version 0.9.17) to produce venn diagrams for comparisons of taxa across cultivars, years, and nutrient media (Russel, 2022). The rare_curve function was used in vegan to produce rarefaction curves for samples. ASVs with less than 5 reads in the generated dataset were removed for beta diversity analysis. The phyloseq package was used to calculate euclidean distances for ASVs and RRPP package (version 1.3.1) was utilized to conduct PERMANOVA analysis on community structure (Collyer & Adams, 2018). Fungal community structure was visualized using principal co-ordinate analysis (PCoA) ordination plots using phyloseq.

## RESULTS

### Seed fragmentation and isolation methods yielded a diverse cohort of isolates

Increasing taxonomic resolution and creating a regionally unique culture library of fungi associated with wheat seed improves our understanding of the role that seed associated fungi play in the developing wheat plant. To examine how cultivar and field seasons impact the wheat seed fungal community, we planted four winter wheat cultivars, Catawba, Hilliard, Shirley, and USG 3640, at Midpines Research Station in Raleigh, NC in 2020 and 2021. Field season 2020 and 2021 were conducted using standard practices with yields ranging from 6.26 lbs to 55.14 lbs per field season across all cultivars. Climatic conditions for 2020 were comparable to previous years for this geographical location. A severe drought occurred in 2021, coinciding with heading and grain filling (Table S1). These conditions required overhead irrigation for field season 2021. We surface disinfested seeds from each cultivar and year to examine the fungi associated with the interior tissue of the seeds, or fungal endophytes. To improve our isolation methods, we experimented with fragment size of macerated seeds, as we observed that split seed isolations often resulted in isolation of a single fast-growing colony (Fig.1) (Bills *et al*., 2004). We selected seed fragments between 177 and 250 µm as they allowed us to capture maximum unique colony morphotypes. We also determined that a dilution factor of 1:100 of wheat seed fragments in sterile water allowed for optimal isolation of unique colonies from the original isolation plate (Fig. 1).

**Figure 1.**
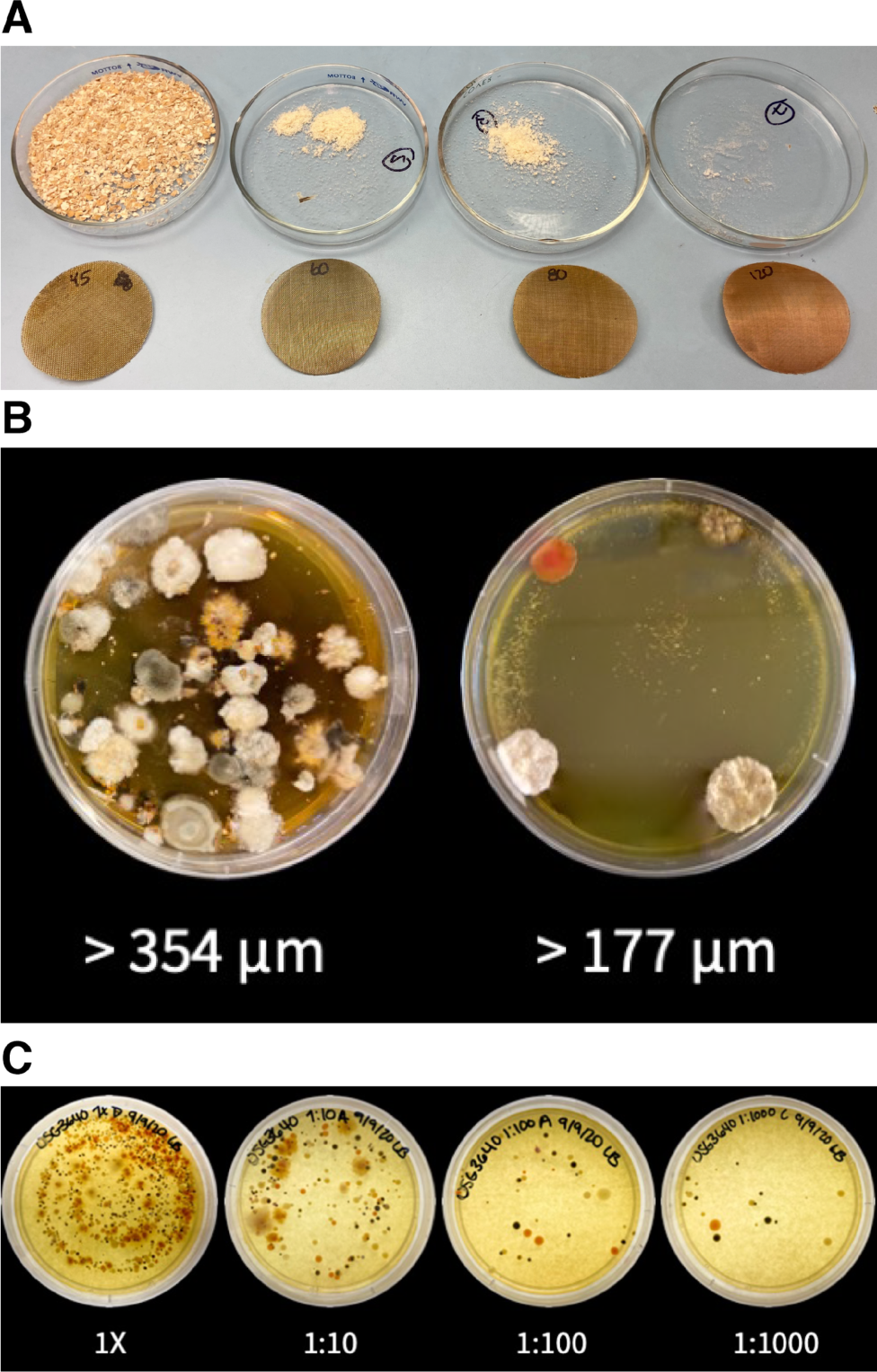
Optimization of fungal isolation from wheat seeds. (A) Particle sizes and associated sieve sizes ranging from 45-120 standard mesh size. (B) Comparison of isolate density in isolation plates with particle filtration sizes > 354 µm and >177 µm. (C) Comparison of dilutions in relation to isolate density, with dilution spanning from 1X to 1:1000.

Three different nutrient media were used to isolate fungi from wheat seed in this study. Four hundred and forty-eight and 197 (total = 645) fungal isolates were obtained from wheat seed sampled in 2020 and 2021, respectively. DG-18 and MYEA yielded a total of 29 and 35 unique fungal morphotypes respectively. Thirty-eight unique morphotypes were obtained from MYEA medium amended with the fungistatic compound cyclosporine (*data not shown*).

### Culture dependent approach yields diverse Ascomycota fungi

To taxonomically classify our isolate collection, we sequenced the ITS1 region and classified sequences using the UNITE database. We recorded a total of 61 unique sequences associated with 21 taxa. The fungal taxa identified were associated with 12 genera, all classified as Ascomycetes that fell within five orders: Capnodiales, Dothideales, Hypocreales, Pleosporales, and Xylariales (Fig. 2). Our intensive culturing technique also allowed us to approximate relative abundance of genera associated by cultivar, year (Fig. 2A) and media (Fig. 2B.). The variation observed suggests that including multiple cultivars, field seasons, and media aids in capturing diverse taxa that may otherwise be omitted. We observed a large share of taxa (N = 11) unique to a single cultivar, while only five taxa occurred across all four cultivars (Fig. S1). The five shared taxa included *Alternaria alstroemeriae*, *Epicoccum layuense*, *Cladosporium* sp., *Parastagonospora poagena*, and *P. forlicesenica*. When examining media types, nine taxa were unique to a single media type, with the largest share attributed to DG18 (N = 5), and fewer unique taxa associated with MYEA (N = 2) and MYEA-CYC (N = 2) (Fig. S1). Our most striking difference was observed by year, with only four taxa shared by the two years examined, comprising of *Alternaria alstroemeriae*, *Alternaria arbusti*, *Pyrenophora nisikadoi*, and *Parastagonospora poagena* (Fig. S1). Our culture dependent efforts generated a broad range of fungal isolates classified within the Ascomycota phylum and illustrated the importance of surveying multiple cultivars with various nutrient media.

**Figure 2.**
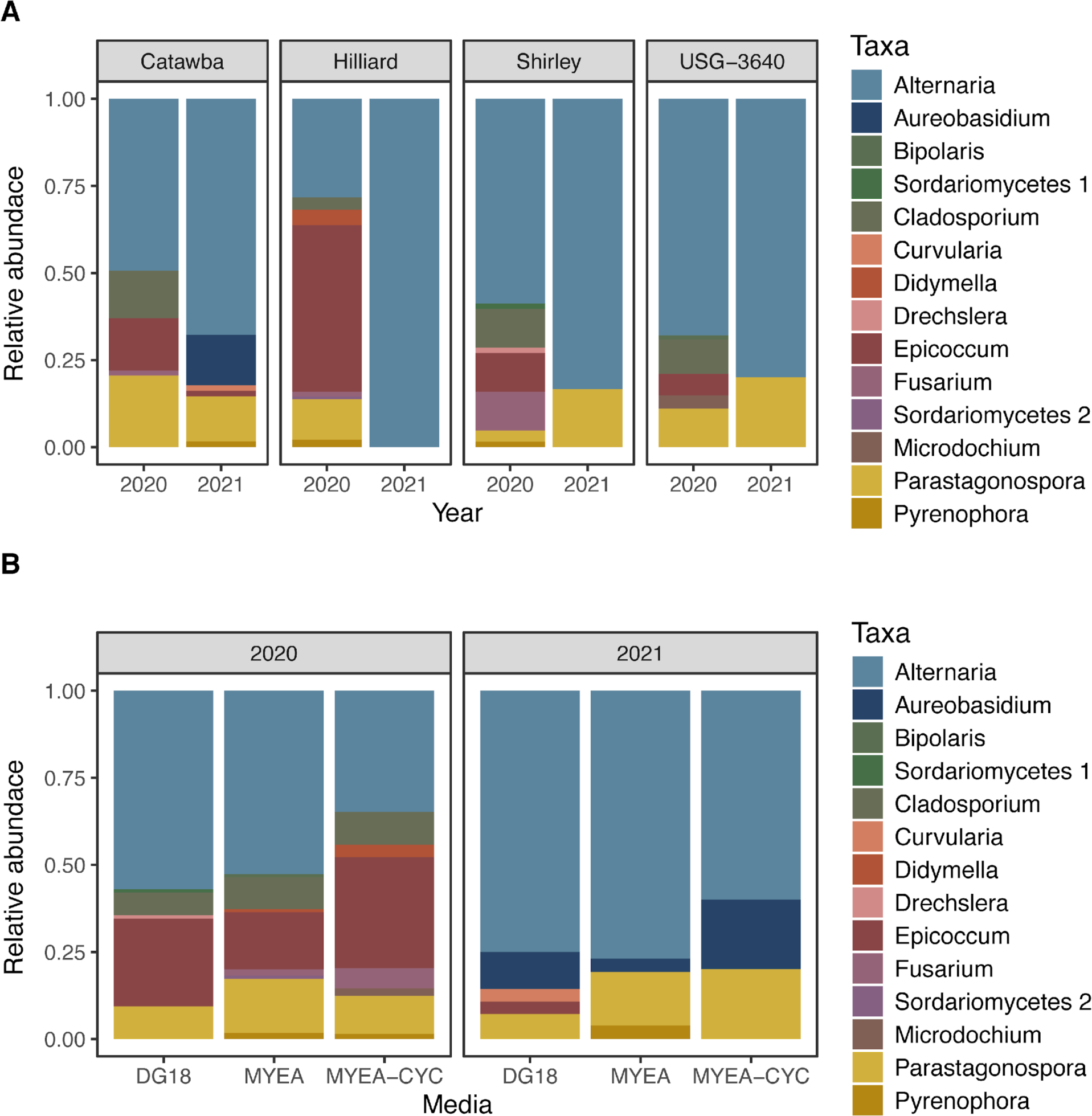
A) Relative abundance of fungal genera by cultivar and year for culture dependent analysis. B) Relative abundance of fungal genera by nutrient media and year for culture dependent analysis.

### Culture independent approach captured a broad diversity of taxa within seeds

A total of 1,670,217 reads were captured with an average of 52,194 reads per sample prior to sequence processing. Quality filtering and standard denoising of reads resulted in an average of 42,240 reads per sample. Forward and reverse reads were merged and chimeric reads were removed, resulting in 1,180,754 total reads with 36,898 reads per sample utilized for further analysis. The number of reads per individual seed sample ranged from 12,180 to 98,659 reads. Sample saturation via a rarefaction curve reached a plateau around 10,000 reads, largely due to the low diversity of the seed mycobiome for individual samples (Fig. S2). After employing Lulu curation to remove spurious ASVs, we detected 78 unique ASVs within our dataset. Fifty unique ASVs were associated with 2020 samples, while 42 unique ASVs were associated with 2021 samples. Only 14 ASVs were detected in both years. The UNITE database was utilized to classify the remaining ASVs, yielding taxa within the Ascomycota, Basidiomycota, and Chytridiomycota, representing 31 distinct genera.

### Wheat seed fungal species richness varied by year

Comprehensive understanding of the diversity of fungal taxa harbored within wheat seeds requires examination of richness and abundance of endophytic fungal taxa. To examine the diversity of taxa associated with wheat seeds, we performed an ANOVA analysis of the observed richness metric for fungal taxa associated with each cultivar and year. Seed samples ranged from 5 to 15 unique ASVs in 2020 and 5 to 11 unique ASVs per seed in 2021. When comparing species richness by cultivar we did not observe significant differences (*P* = 0.770), indicating stability of the fungal seed endophytic community. However, there was a decrease in observed richness by year, with 2021 exhibiting lower richness compared to 2020 (*P* = 0.032) (Fig. 3). This decrease in richness mirrors the decrease in observed morphotypes that we recorded for our culture dependent results. To examine taxonomic consistency between the two years, we classified the fourteen shared ASVs, and found co-occurrence for both years of *Alternaria*, *Aureobasidium*, *Bullera*, *Cladosporium*, *Dioszegia*, *Epicoccum, Monographella*, *Neoascochyta*, *Papiliotrema*, *Parastagonospora*, and *Stemphylium*. Notably, only *Alternaria*, *Epicoccum*, and *Parastagonospora* were captured in both years for the culture dependent study. *Alternaria, Cladosporium, Epicoccum, Monographella,* and *Parastagonospora* were present in all cultivars based on metabarcoding analysis, similar to the culture dependent analysis with the exception of *Monographella* (Figs. 2 and 4).

**Figure 3.**
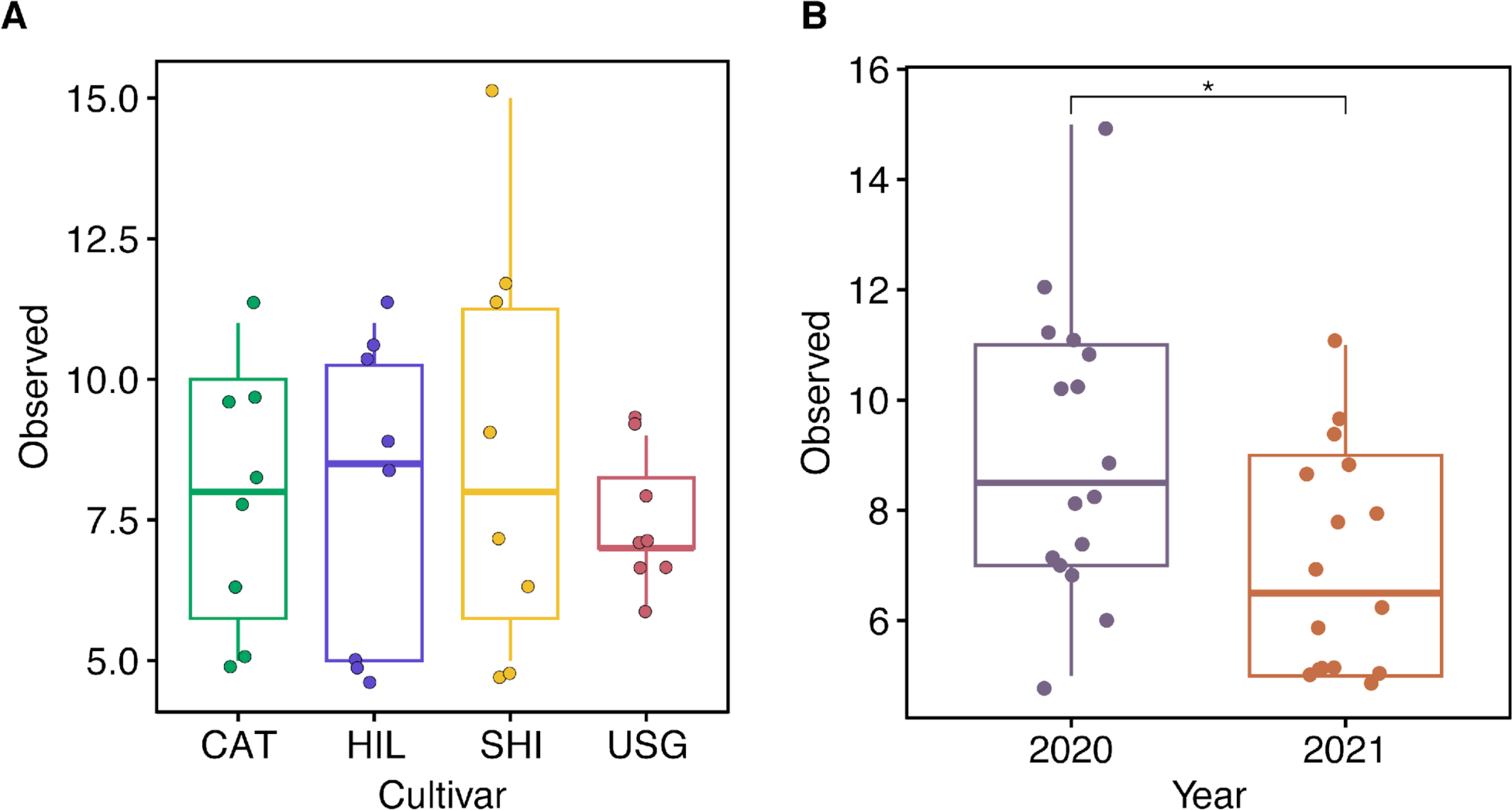
Alpha diversity of fungi associated with wheat seeds estimated by Observed richness by A) Cultivar and B) Year. For each box plot, the whiskers mark the minimum and maximum vlaues, while the top and bottom lines of each box represent third and first quartiles. The center line represents the median value. Analysis of variance (ANOVA) and Tukey’s honestly significant difference (HSD) were utilized to examine significant variation among cultivars and years (*P* < 0.05).

**Figure 4.**
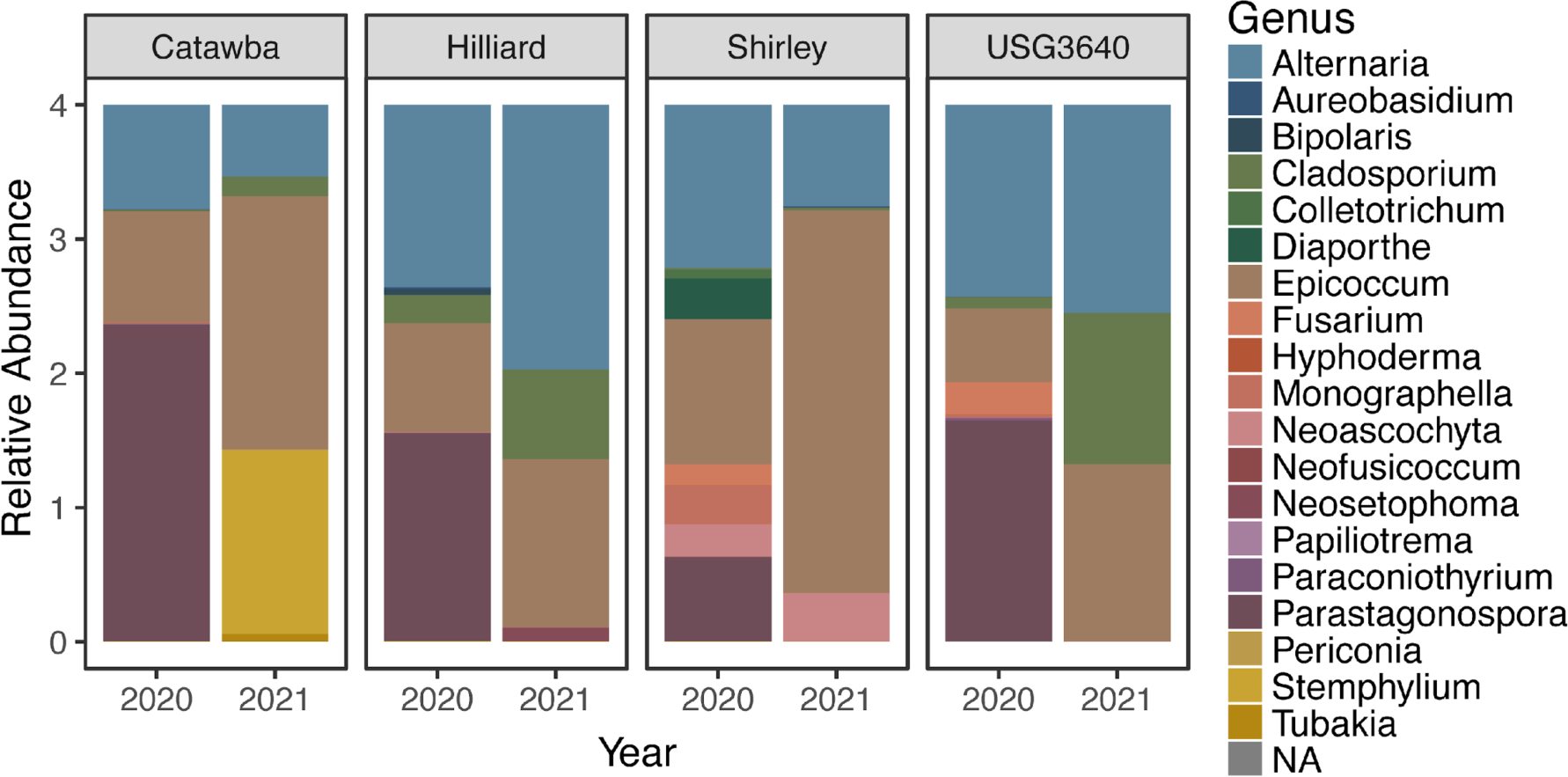
Relative abundance of fungal genera associated with thirty most abundant ASVs by cultivar and year for culture independent analysis.

When examining the abundance of the most frequent genera associated with seeds, we noted that *Alternaria* and *Epicoccum* played substantial and consistent roles in shaping the seed mycobiome for all cultivars and years in both culture independent and culture dependent anlayses (Figs. 2 and 4). *Parastagonospora* was abundant for all cultivars in the 2020 season, but largely absent in the 2021 season based on metabarcoding, in contrast to our culture dependent approach, which indicated that *Parastagonospora* was equally abundant across years (Figs. 2 and 4). *Cladosporium* was present in both Hilliard and USG3640 and exhibited a higher relative abundance in 2021 compared to 2020, whereas in our culture dependent survey *Cladosporium* was mostly captured within our 2020 samples (Figs. 2 and 4). Other genera uniquely associated with a single cultivar, such as *Neoascochyta’s* association with Shirley, which was not captured in our culture dependent analysis. To a lesser extent, other taxa were unique either to a few cultivars, such as *Fusarium’s* affiliation with Shirley and USG 3640 in 2020, which was only partially confirmed by *Fusarium’s* association with Catawba, Hilliard, and Shirley in 2020 for culture dependent results (Figs. 2 and 4). *Stemphylium’s* association with Catawba in 2021 was not captured by culture dependent analysis. Overall richness decreased in 2021 for both culture dependent and independent approaches, with a substantial shift in taxa abundance across both years.

### Compositional analysis of the wheat seed mycobiome lacks structure

Understanding factors that contribute to the composition of the fungal community structure is key to understanding the rules of seed mycobiome assembly. To examine if cultivar or year structured the wheat seed fungal community, we conducted a PERMANOVA analysis on our culture independent data set, using euclidean distances. Neither cultivar (*P* = 0.35) or year (*P* =0.26) had a significant effect on the structure of wheat seed mycobiome. When visualizing the fungal community structure with euclidean distances using a PCoA, the first two axes accounted for 22.8% and 17.7% of the variance observed, respectively (Fig. S3). Visually there was no separation by cultivar or year, consistent with our PERMANOVA analysis. Although we observed that year played a role in influencing species richness, cultivar and year did not appear to account for a significant portion of variance within the fungal community structure for our culture independent analysis. To visualize fungal genera associated with culture-independent investigation of the wheat seed mycobiome, we generated a heat tree. The size and color of the nodes indicate occurrence of taxa matching to our 78 ASVs (Fig. 5). We detected Basidiomycete taxa in our culture independent approach; however they were absent from our culture dependent analysis. Basidiomycete taxa detected within seed samples included *Papiliotrema*, *Dioszegia*, and *Filobasidium*. In addition, *Nigrospora*, *Colletotrichum*, *Tubakia*, *Monographella* and *Diaporthe*, all Sordariomycetes, were only captured with our culture independent methods. Within the Dothideomycetes, *Neofusicoccum*, *Periconia*, *Paraconiothyrium*, *Phaesophaeria*, *Neoascochyta* and *Neosetophoma*, were all unique to our culture independent methods (Fig. 5). When examining fungal taxa unique to our culture dependent methods, the list of genera include *Bipolaris*, *Curvularia*, *Didymella*, *Dreschlera*, and *Microdochium*. The most abundant taxa associated with our culture independent methods; *Alternaria*, *Epicoccum*, *Cladosporium*, and *Parastagonospora*, were consistently isolated with culture dependent methods. Taken together, our results highlight the importance of combining culture independent and dependent methods to capture and establish a diverse endophytic fungal catalog associated with the wheat seed.

**Figure 5.**
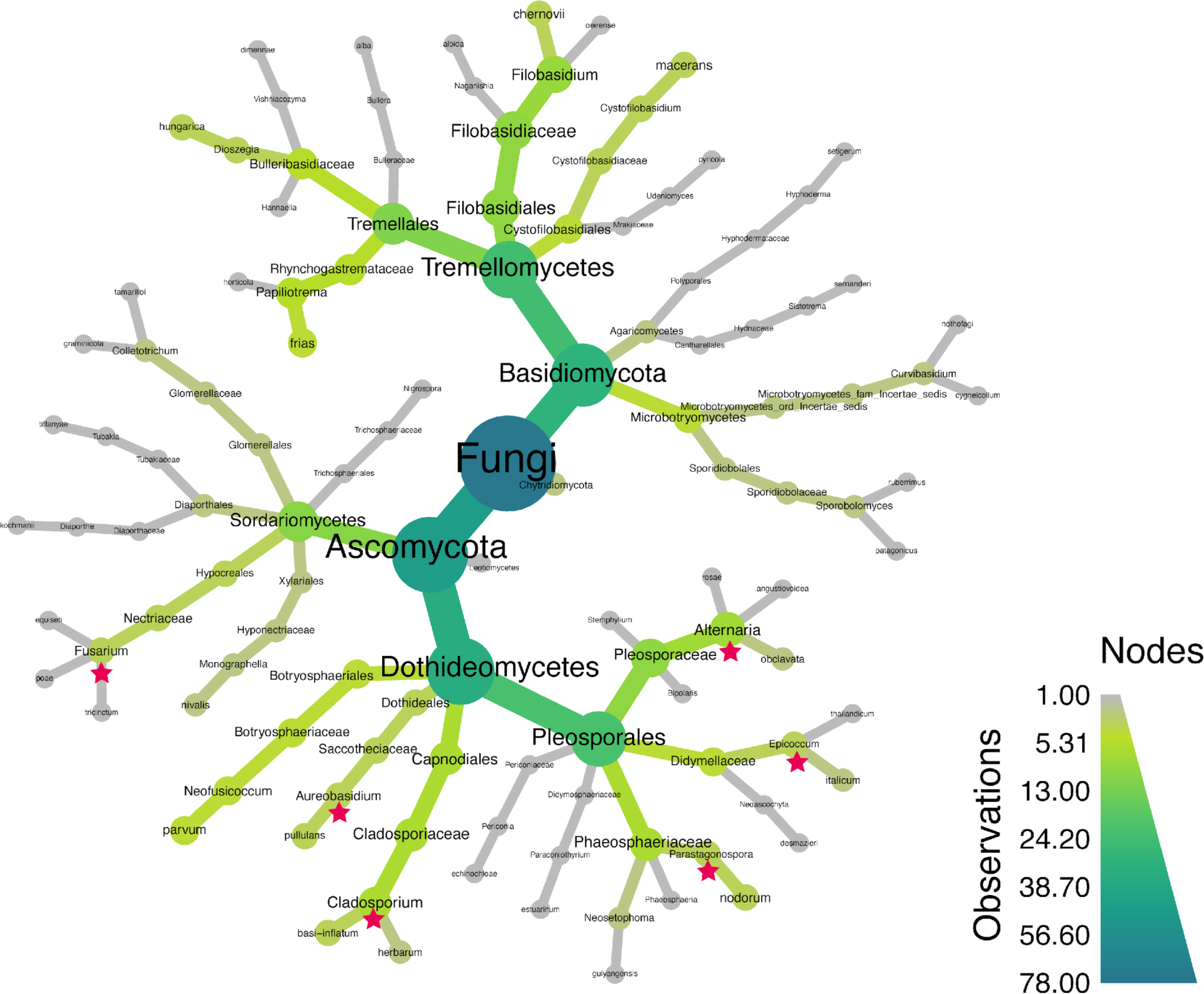
Heat tree of fungal taxa associated with wheat seeds in culture independent survey. Color of the individual branches and nodes indicates number of observations of a particular taxa within the dataset of 78 ASVs. Color of genera names indicates if they are found in both culture independent and dependent surveys, or only in culture dependent surveys.

## DISCUSSION

In this study, we utilized a synergistic culture independent and culture dependent experimental approach with the goal of capturing comprehensive diversity of the wheat seed mycobiome and to curate a culture library of regionally unique fungal isolates. We identified a substantial number of taxa at the species level, across several taxonomic classes, and within the phylum Ascomycota by implementing a culture dependent approach. Amplicon sequencing of the ITS1 rDNA region allowed for detection of a broad group of diverse taxa across several phyla. The variation that we observed in our fungal communities by year, cultivar, and approach highlights the success of our refined methods to capture diverse taxa and also pinpoint areas of improvement to expand our culture library. This dual methods approach has been previously deployed in wheat to isolate fungi from two wheat cultivars to test the primary symbiont hypothesis in the Pacific Northwest and in switchgrass to identify endophytic fungal isolates that play a role in drought tolerance (Giauque & Hawkes, 2013; Ridout *et al*., 2019). Our culture dependent approach exemplified how variation in fragment size, dilutions, and media types can yield diverse taxa. In contrast with previous studies, we utilized three distinct nutrient media to mimic the internal seed environment (Larran *et al*., 2007) and facilitate isolation of slow growing fungi by incorporating cyclosporine, originally developed to isolate rare endophytic fungal taxa from grass and deciduous forest litter (Bills *et al*., 2004; Pirttilä & S, 2011). The most commonly isolated taxa from wheat seeds were, *Alternaria*, *Cladosporium*, *Epicoccum*, and *Parastagonospora* sampled from a variety of wheat cultivars and production regions over the last three decades (Sieber *et al*., 1988; Larran *et al*., 2002; Comby *et al*., 2016; Ridout *et al*., 2019). Less commonly isolated genera that we identified in our study included *Bipolaris*, *Didymella*, *Fusarium*, *Microdochium*, *Aureobasidium*, and *Dreschlera* (Sieber *et al*., 1988; Larran *et al*., 2007; Comby *et al*., 2016). The Ascomycete *Curvularia* isolated in 2021 from Catawba seeds represents a first reported instance of *Curvularia* isolated from wheat seeds to our knowledge. However, *Curvularia* has been previously isolated from wheat glumes and leaves, suggesting a potential source of this fungal endophyte (Larran *et al*., 2007). Our culture dependent experimental approach yielded a diverse group of Ascomycota taxa, consistent with previous surveys of wheat seeds of five wheat cultivars from Argentina (Larran *et al*., 2007). Notably, our culture library lacked representation from members of the Basidiomycota phylum, which are often sampled at a low frequency in culture-dependent surveys (Sieber *et al*., 1988; Comby *et al*., 2016; Solanki *et al*., 2019). When examining the fungal communities of wheat seeds sourced from cultivars with varying susceptibility to Fusarium head blight, Comby et al. (2016) isolated only two Basidiomycete taxa, *Trametes hirsuta* and *Rhodosporidium kratochvilovae*, from wheat seeds using a standard malt agar nutrient medium. The recalcitrant isolation of Basidiomycetes from wheat seeds was first reported in Sieber *et al*. (1988) exhaustive isolation approach which yielded nearly 27,000 isolates, with only 2% corresponding to Basidiomycetes, specifically *Rhizoctonia solani*. The fungal taxa isolated in our culture dependent approach reflect global trends of wheat seed associated fungi, encompassing a contemporary and regionally specific endophyte collection.

Variation observed in our culture dependent efforts across years may potentially reflect the late season drought experienced in central NC in 2021, which required periodic overhead irrigation of plots with pond water. A sharp decline in isolates was observed in 2021, and corresponded with a decrease in species richness in 2021 of metabarcoded seed samples. We also observed that wheat cultivars with the lowest species richness in 2021 for culture dependent analysis exhibited similar decreases in culture independent analysis, highlighting that our isolation methods captured overall trends in the wheat seed mycobiome. Notably, the most abundant taxa in 2021 were largely restricted to *Alternaria*, *Aureobasidium*, and *Parastagonospora*. Official disease ratings were not conducted in our plots, however *Parastagonospora* is well known as a wheat pathogen within North Carolina field sites and can be transmitted as a seedborne pathogen in the eastern US (Bennett *et al*., 2007). *Parastagonospora* favors high humidity accumulated in kernels, which can be attributed to overhead irrigation. We hypothesize that overhead irrigation applied during late season 2021 favored colonization of *Parastagonospora* and promoted excessive growth of *Alternaria* (Bockus *et al*., 2010; Somma *et al*., 2019). Our culture-dependent efforts isolated taxa classified as pathogenic or may contribute to grain spoilage, however, pathogenicity assays must be conducted to confirm their lifestyle at the individual isolate level (Bockus *et al*., 2010; DeMers, 2022).

The seed environment represents a unique niche that rapidly changes over time to become a water limited environment with shifting carbohydrate sources (Bewley & Black, 1994). These conditions have been hypothesized to influence the occurrence of fungal taxa associated with seeds in addition to pathogens acting latently as the seeds serve as a source for transmission among generations (Nelson, 2018). While seed transmission of pathogenic taxa remains a concern for producers, seeds also harbor non-pathogenic fungal taxa that can be transmitted among generations (Vujanovic *et al*., 2019). Recent studies have noted the important roles that seed sourced fungal endophytes can play in developing seedlings, with seed associated *Fusarium* and *Alternaria* detected in early germination and seedling stages of wheat, rice, barley, soybean, common bean, and oaks (Abdelfattah *et al*., 2021; Johnston-Monje *et al*., 2021). The transmission of non-pathogenic and possibly beneficial fungi among generations warrants future examination of isolate-level genomic tracking between seeds and seedlings (Mitter *et al*., 2017).

Our metabarcoding results largely mirrored trends observed in our culture dependent results, with *Alternaria*, *Cladosporium*, *Epicoccum* and *Parastagonospora* dominating as the most abundant taxa within the majority of our samples. These results are consistent with other wheat seed endophytic metabarcoding surveys (Hertz *et al*., 2016; Latz *et al*., 2021). Notably, other surveys of the wheat seed mycobiome observed higher occurrence and abundance of *Fusarium* species while we observed only sporadic associations among the cultivars examined in this study (Nicolaisen *et al*., 2014; Bakker & McCormick, 2019; Latz *et al*., 2021b). When examining genera unique to a single cultivar and year, *Stemphylium*, implicated as a causal agent of Sooty Head Molds, was abundant within Catawba seeds in 2021 (Bockus *et al*., 2010). Fungal genera not detected within our culture-dependent assay included *Monographella*, *Neofusicoccum*, and *Neoascochyta*, all of which were detected using metabarcoding methods in other studies but rarely isolated in culture (Solanki *et al*., 2019; Latz *et al*., 2021). Our metabarcoding analysis revealed an increased occurrence of fungal taxa in the Basidiomycota, largely representing yeasts associated with wheat seeds such as *Dioszegia*, *Filobasidium*, *Papiliotrema*, and *Sporobolomyces* (Nicolaisen *et al*., 2014; Grudzinska-Sterno *et al*., 2016). *Sporobolomyces* has been suggested to be an early colonizer during wheat seed development and is hypothesized to exhibit a competitive interaction with *Alternaria infectoria* (Hertz *et al*., 2016). While we cannot confirm that *Alternaria infectoria* was present in wheat seeds, we hypothesize that *Sporobolomyces* co-occurs with other *Alternaria* species in our study. The Basidiomycetes identified with our culture independent approach revealed a set of taxa that lacked co-occurrence with our culture dependent approach, indicating that selective nutrient media for filamentous and yeast forms in this phylum may increase the diversity of taxa observed when coupled with fungal community metabarcoding methods. When examining taxa that were only present in our culture dependent approach, several taxa in the family *Pleosporaceae* such as *Bipolaris*, *Curvularia*, *Dreschlera*, and other taxa including *Didymella*, and *Microdochium* were absent in our metabarcoding analysis. We note that our culture dependent approach was more robust as it incorporated a pool of seeds from each cultivar for each nutrient media, while our culture independent approach focused on a single seed metabarcoding survey. While examining the fungal richness and communities of individual seeds allows for seed-level analysis, this method may have led to lower species richness observed in our metabarcoding approach and presents a major limitation of our study.

Our wheat seed mycobiome analysis revealed a consistent fungal community that remains stable across cultivars and two field seasons. We hypothesize that minor variation in species richness across years and lack of variation in community structure was largely due to complexity and the stochasticity of the wheat seed mycobiome. Previous studies examining traits such as wheat cultivar relatedness and disease susceptibility concluded that neither factor appears to influence the composition or structure of the wheat seed mycobiome (Comby *et al*., 2016; Latz *et al*., 2021). Environmental factors such as whether plants were grown in a field or in a greenhouse can account for a large proportion of variation within the wheat seed mycobiome, as well as stages of wheat grain development (Hertz *et al*., 2016; Latz *et al*., 2021). Other metabarcoding surveys of the wheat seed mycobiome over field seasons provided experimental evidence for increased species richness (Nicolaisen *et al*., 2014). These results are similar to what we observed in 2020, which may have been attributed to a growing season with more rainfall that potentially increased the occurrence of rare taxa in our what seed samples. Within the grass family *Poaceae*, the dominant associations of seed-transmitted *Epichloë* fungi are well documented in fescue (Schardl *et al*., 2004). When examining trends observed in other seed mycobiomes for other plant genera, the culturable fungal taxa isolated from single seeds tends to be limited to one or two taxa, and in some studies none (Newcombe *et al*., 2018). This observation has led to the hypothesis that the seed niche likely selects for a limited number of taxa capable of bypassing the external anatomical barriers of the plant and are able to adapt and survive changes in the seed environment (Newcombe *et al*., 2018). In our study, we observed that climatic conditions may affect species richness determining the pool of the possible seed associated taxa. While physical barriers may reduce the candidate cohort of the seed associated fungal community, the extent of the influence that plant immunity plays in determining the seed mycobiome remains poorly understood for non-mycorrhizal fungi (Jones & Dangl, 2006; Vannette, 2020). Taken together, culture dependent and independent approaches can expand our knowledge and understanding of the wheat seed mycobiome. The culture library generated in this study provides us with an opportunity to conduct future research to improve taxonomic and genomic resolution of fungal isolates and examine the functional role that they play in determining adult plant outcomes (Noel *et al*., 2022).

## ACKNOWLEDGEMENTS

We would like to thank Myron Fountain and Charlie Glover (USDA-ARS Raleigh), who provided seed for Catawba, Shirley and Hilliard and designed field plot experiments, carried out field experiments, and provided vast technical knowledge and assistance throughout the project. We are grateful to Gina Brown-Guedira USDA-ARS Raleigh, who provided seed for USG 3640. Thanks to our collaborators in INTERACT consortium who provided insight on methods used in this study, especially Jens Frisvad. Gerald Bills provided additional insight into isolation methods with particle sieving and addition of cyclosporine. We are indebted to Brian Gilger who provided us with cyclosporine. We thank Christine Hawkes for her guidance and expertise in metabarcoding analysis. We acknowledge the computing resources provided by North Carolina State University High Performance Computing Services Core Facility (RRID:SCR_022168). We thank Lisa Lowe for her guidance and support for using the computing services efficiently and Parker Ingraham for her invaluable help in culture isolation and skillful DNA extractions from isolates and seeds. This work was funded by NovoNordisk Fonden Grant #NNF19SA0059360.

---

**Table S1.** Monthly weather data for Midpines research station in Raleigh, NC. Field season is indicated as 2020 or 2021 for harvest year. Maximum air temperature, average air temperature, total precipitation, and average relative humidity are presented below.

| **Field season** | **Month** | **Average Maximum Air Temperature (C)** | **Average Air Temperature (C)** | **Total Precipitation (mm)** | **Average Relative Humidity (%)** |
| --- | --- | --- | --- | --- | --- |
| 2020 | 2019-10 | 18.7 | 25 | 92.96 | 72.16 |
| 2020 | 2019-11 | 14.4 | 8.6 | 87.12 | 68.48 |
| 2020 | 2019-12 | 13.8 | 8.5 | 103.89 | 71.97 |
| 2020 | 2020-01 | 13.2 | 8.3 | 138.18 | 68.49 |
| 2020 | 2020-02 | 13.9 | 9 | 153.16 | 69.95 |
| 2020 | 2020-03 | 18.6 | 13.8 | 79.25 | 67.54 |
| 2020 | 2020-04 | 21.2 | 15.1 | 102.36 | 61.95 |
| 2020 | 2020-05 | 23.3 | 18 | 126.75 | 69.82 |
| 2021 | 2020-10 | 17.4 | 23.3 | 70.61 | 77.6 |
| 2021 | 2020-11 | 19.3 | 13.2 | 173.74 | 73 |
| 2021 | 2020-12 | 11.8 | 6.1 | 137.16 | 70.34 |
| 2021 | 2021-01 | 9.7 | 4.9 | 155.7 | 70.75 |
| 2021 | 2021-02 | 10.7 | 5.7 | 178.05 | 70.09 |
| 2021 | 2021-03 | 18.2 | 12.5 | 64.52 | 58.5 |
| 2021 | 2021-04 | 22.6 | 15.9 | 16 | 54.42 |
| 2021 | 2021-05 | 26.1 | 19.2 | 50.29 | 62.14 |

**Figure S1.**
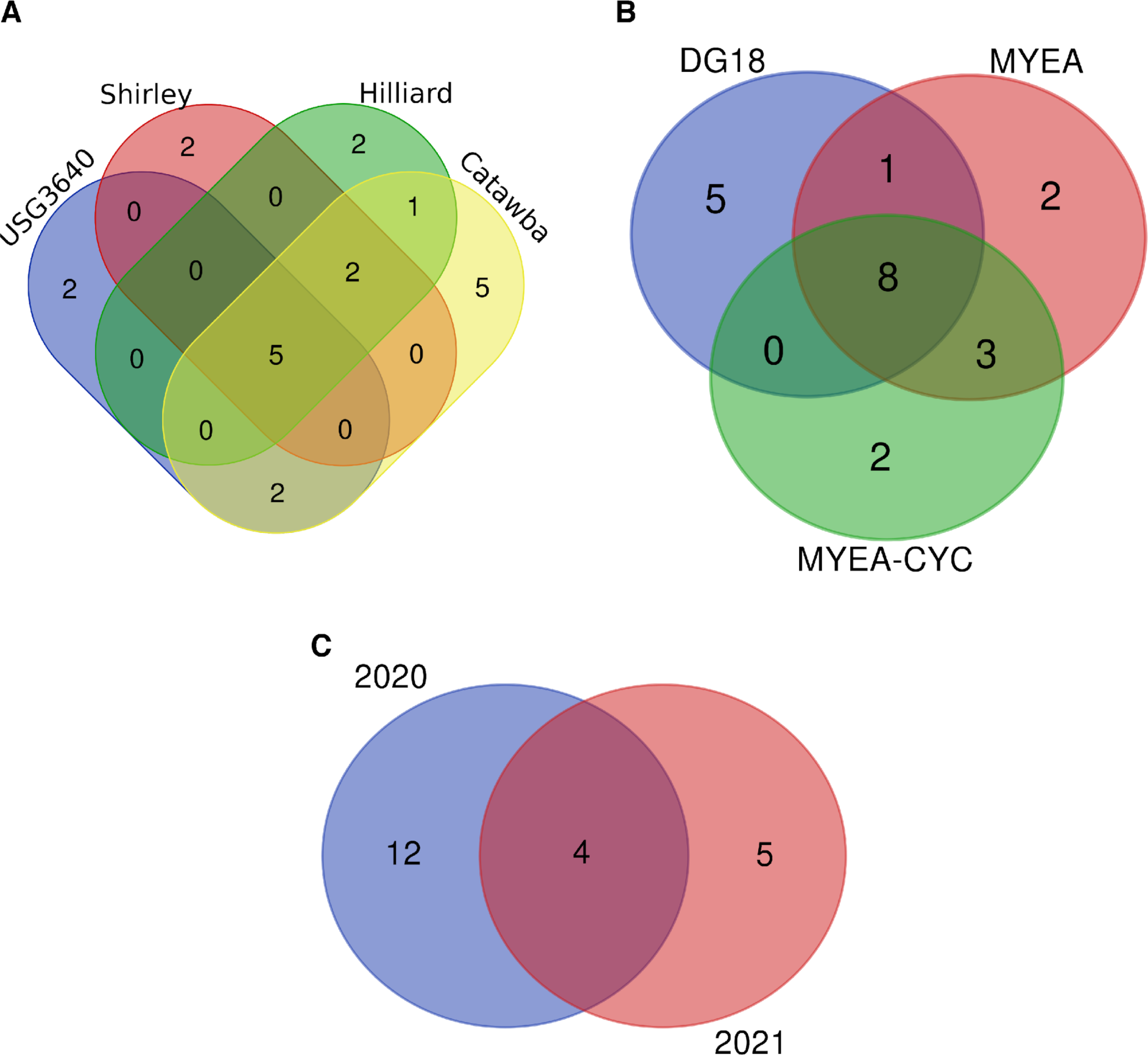
Venn diagrams indicating overlap at the species level among A) cultivars, B) nutrient media types, C) years for culture dependent approach.

**Figure S2.**
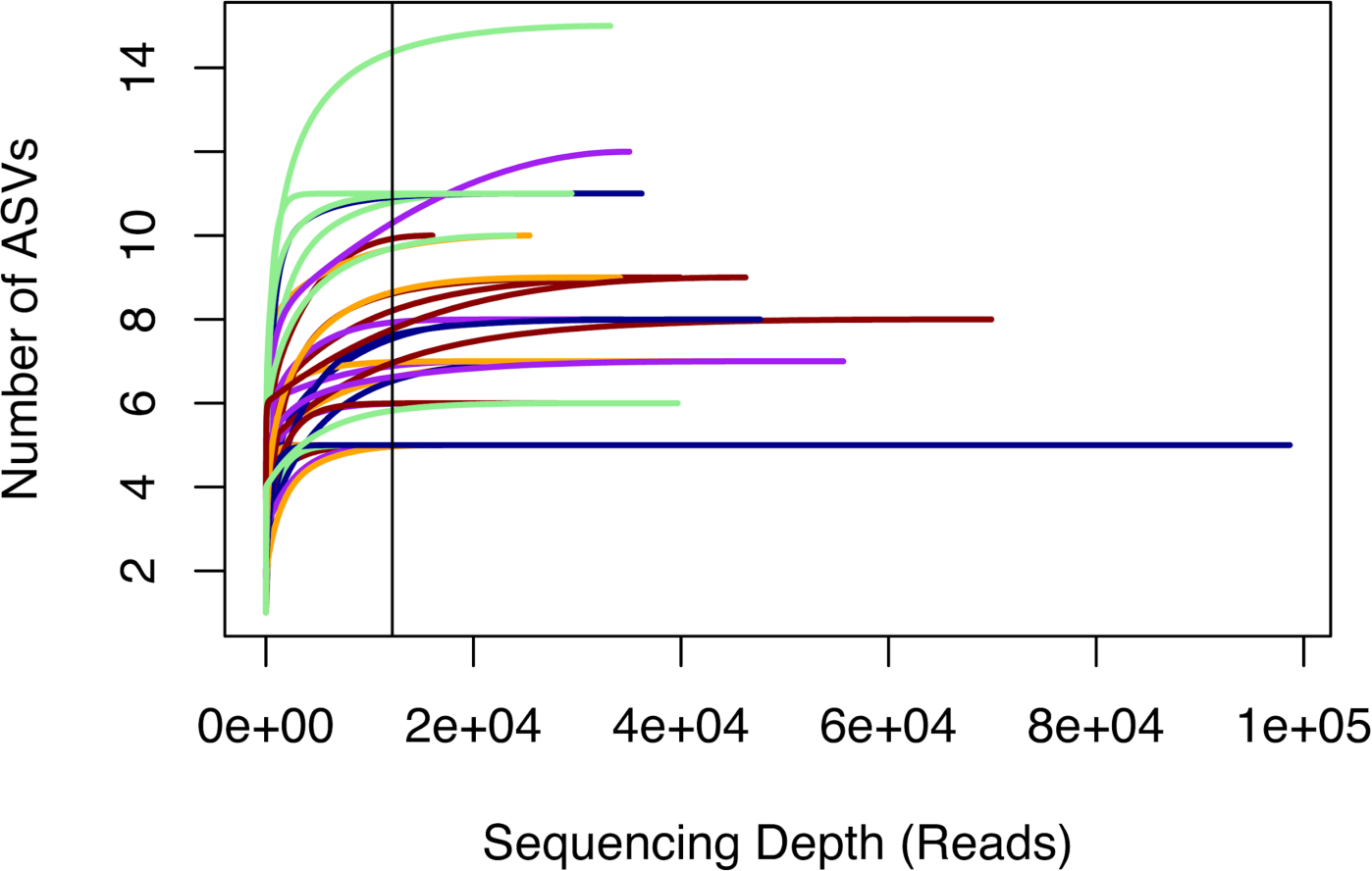
Sample rarefaction curve of samples by their read depth for metabarcoding approach. Vertical line indicates minimum read depth of 12,180

**Figure S3.**
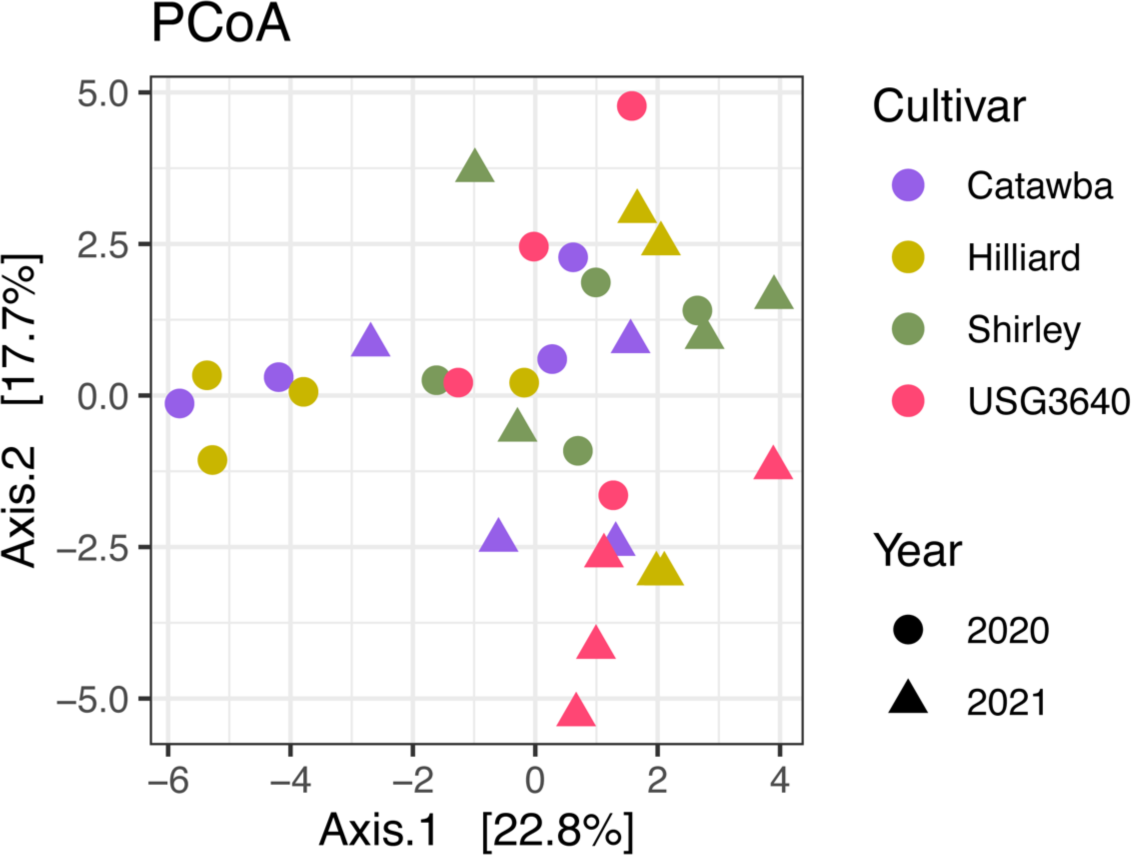
Principal Co-ordinates Analysis plot based on Euclidean Distances to examine structure of the wheat seed mycobiome by cultivar and year.

## LITERATURE CITED

Abarenkov K, Henrik Nilsson R, Larsson K-H, Alexander IJ, Eberhardt U, Erland S, Høiland K, Kjøller R, Larsson E, Pennanen T, et al.2010. The UNITE database for molecular identification of fungi – recent updates and future perspectives. New Phytologist 186: 281–285.

Abdelfattah A, Wisniewski M, Schena L, Tack AJM. 2021. Experimental evidence of microbial inheritance in plants and transmission routes from seed to phyllosphere and root. Environmental Microbiology 23: 2199–2214.

Agarwal VK, Sinclair JB. 2014. Principles of Seed Pathology. Boca Raton: CRC Press.

Anguita-Maeso M, Olivares-García C, Haro C, Imperial J, Navas-Cortés JA, Landa BB. 2020. Culture-Dependent and Culture-Independent Characterization of the Olive Xylem Microbiota: Effect of Sap Extraction Methods. Frontiers in Plant Science 10.

Bakker MG, McCormick SP. 2019. Microbial Correlates of Fusarium Load and Deoxynivalenol Content in Individual Wheat Kernels. Phytopathology 109: 993–1002.

Bennett RS, Milgroom MG, Sainudiin R, Cunfer BM, Bergstrom GC. 2007. Relative Contribution of Seed-Transmitted Inoculum to Foliar Populations of *Phaeosphaeria nodorum*. Phytopathology 97: 584–591.

Bergna A, Cernava T, Rändler M, Grosch R, Zachow C, Berg G. 2018. Tomato Seeds Preferably Transmit Plant Beneficial Endophytes. Phytobiomes Journal 2: 183–193.

Bewley JD, Black M. 1994. Seed Development and Maturation. In: Bewley JD, Black M, eds. Seeds: Physiology of Development and Germination. Boston, MA: Springer US, 35–115.

Bills GF, Christensen M, Powell M, Thorn G. 2004. 13 - SAPROBIC SOIL FUNGI. In: Mueller GM, Bills GF, Foster MS, eds. Biodiversity of Fungi. Burlington: Academic Press, 271–302.

Bintarti AF, Sulesky-Grieb A, Stopnisek N, Shade A. 2022. Endophytic Microbiome Variation Among Single Plant Seeds. Phytobiomes Journal 6: 45–55.

Bockus WW, Bowden RL, Hunger RM, Morrill WL, Murray TD, Smiley RW. 2010. Diseases Caused by Fungi and Fungus-Like Organisms. In: Diseases and Pests Compendium Series. Compendium of Wheat Diseases and Pests, Third Edition. The American Phytopathological Society, 15–86.

Calderini DF, Castillo FM, Arenas-M A, Molero G, Reynolds MP, Craze M, Bowden S, Milner MJ, Wallington EJ, Dowle A, et al.2021. Overcoming the trade-off between grain weight and number in wheat by the ectopic expression of expansin in developing seeds leads to increased yield potential. New Phytologist 230: 629–640.

Callahan BJ, Grinevich D, Thakur S, Balamotis MA, Yehezkel TB. 2021. Ultra-accurate microbial amplicon sequencing with synthetic long reads. Microbiome 9: 130.

Callahan BJ, McMurdie PJ, Rosen MJ, Han AW, Johnson AJA, Holmes SP. 2016. DADA2: High-resolution sample inference from Illumina amplicon data. Nature Methods 13: 581–583.

Choudhury RA, Garrett KA, Klosterman SJ, Subbarao KV, McRoberts N. 2017. A Framework for Optimizing Phytosanitary Thresholds in Seed Systems. Phytopathology 107: 1219–1228.

Christensen CM. 1957. Deterioration of Stored Grains by Fungi. Botanical Review 23: 108–134.

Collyer ML, Adams DC. 2018. RRPP: An r package for fitting linear models to high-dimensional data using residual randomization. Methods in Ecology and Evolution 9: 1772– 1779.

Comby M, Lacoste S, Baillieul F, Profizi C, Dupont J. 2016. Spatial and Temporal Variation of Cultivable Communities of Co-occurring Endophytes and Pathogens in Wheat. Frontiers in Microbiology 7.

DeMers M. 2022. *Alternaria alternata* as endophyte and pathogen. *Microbiology (Reading*, England) 168: 001153.

Díaz Herrera S, Grossi C, Zawoznik M, Groppa MD. 2016. Wheat seeds harbour bacterial endophytes with potential as plant growth promoters and biocontrol agents of *Fusarium graminearum*. Microbiological Research 186–187: 37–43.

Dissanayake AJ, Purahong W, Wubet T, Hyde KD, Zhang W, Xu H, Zhang G, Fu C, Liu M, Xing Q, et al.2018. Direct comparison of culture-dependent and culture-independent molecular approaches reveal the diversity of fungal endophytic communities in stems of grapevine (Vitis vinifera). Fungal Diversity 90: 85–107.

Flannigan B. 1974. Distribution of seed-borne micro-organisms in naked barley and wheat before harvest. Transactions of the British Mycological Society 62: 51–58.

Foster Z, Sharpton T, Grünwald N. 2017. Metacoder: An R package for visualization and manipulation of community taxonomic diversity data. PLOS Computational Biology 13: 1–15.

Frøslev TG, Kjøller R, Bruun HH, Ejrnæs R, Brunbjerg AK, Pietroni C, Hansen AJ. 2017. Algorithm for post-clustering curation of DNA amplicon data yields reliable biodiversity estimates. Nature Communications 8: 1188.

Gagic M, Faville MJ, Zhang W, Forester NT, Rolston MP, Johnson RD, Ganesh S, Koolaard JP, Easton HS, Hudson D, et al.2018. Seed Transmission of Epichloë Endophytes in Lolium perenne Is Heavily Influenced by Host Genetics. Frontiers in Plant Science 9: 1580.

Gdanetz K, Noel Z, Trail F. 2021. Influence of Plant Host and Organ, Management Strategy, and Spore Traits on Microbiome Composition. Phytobiomes Journal 5: 202–219.

Gdanetz K, Trail F. 2017. The Wheat Microbiome Under Four Management Strategies, and Potential for Endophytes in Disease Protection. Phytobiomes Journal 1: 158–168.

Giauque H, Hawkes CV. 2013. Climate affects symbiotic fungal endophyte diversity and performance. American Journal of Botany 100: 1435–1444.

Gitaitis R, Walcott R. 2007. The epidemiology and management of seedborne bacterial diseases. Annual Review of Phytopathology 45: 371–397.

Griffey C, Malla S, Brooks W, Seago J, Christopher A, Thomason W, Pitman R, Markham R, Vaughn M, Dunaway D, et al.2020. Registration of ‘Hilliard’ wheat. Journal of Plant Registrations 14: 406–417.

Griffey CA, Thomason WE, Pitman RM, Beahm BR, Paling JJ, Chen J, Gundrum PG, Fanelli JK, Kenner JC, Dunaway DW, et al.2010. Registration of ‘Shirley’ Wheat. Journal of Plant Registrations 4: 38–43.

Grudzinska-Sterno M, Yuen J, Stenlid J, Djurle A. 2016. Fungal communities in organically grown winter wheat affected by plant organ and development stage. European Journal of Plant Pathology 146: 401–417.

Hardoim PR, van Overbeek LS, Berg G, Pirttilä AM, Compant S, Campisano A, Döring M, Sessitsch A. 2015. The Hidden World within Plants: Ecological and Evolutionary Considerations for Defining Functioning of Microbial Endophytes. Microbiology and Molecular Biology Reviews 79: 293–320.

Hassani MA, Özkurt E, Franzenburg S, Stukenbrock EH. 2020. Ecological Assembly Processes of the Bacterial and Fungal Microbiota of Wild and Domesticated Wheat Species. Phytobiomes Journal 4: 217–224.

Hertz M, Jensen IR, Jensen LØ, Thomsen SN, Winde J, Dueholm MS, Sørensen LH, Wollenberg RD, Sørensen HO, Sondergaard TE, et al. 2016. The fungal community changes over time in developing wheat heads. International Journal of Food Microbiology 222: 30–39.

Hone H, Mann R, Yang G, Kaur J, Tannenbaum I, Li T, Spangenberg G, Sawbridge T. 2020. Drought Tolerant Wheat Varieties Enrich Seed Microbiomes For Beneficial Microbes Under Drought Conditions. In Review.

Johnston-Monje D, Gutiérrez JP, Lopez-Lavalle LAB. 2021. Seed-Transmitted Bacteria and Fungi Dominate Juvenile Plant Microbiomes. Frontiers in Microbiology 12.

Jones JDG, Dangl JL. 2006. The plant immune system. Nature 444: 323–329.

Karlsson I, Friberg H, Kolseth A-K, Steinberg C, Persson P. 2017. Organic farming increases richness of fungal taxa in the wheat phyllosphere. Molecular Ecology 26: 3424–3436.

Kavamura VN, Hayat R, Clark IM, Rossmann M, Mendes R, Hirsch PR, Mauchline TH. 2018. Inorganic Nitrogen Application Affects Both Taxonomical and Predicted Functional Structure of Wheat Rhizosphere Bacterial Communities. Frontiers in Microbiology 9.

Kavamura VN, Robinson RJ, Hayat R, Clark IM, Hughes D, Rossmann M, Hirsch PR, Mendes R, Mauchline TH. 2019. Land Management and Microbial Seed Load Effect on Rhizosphere and Endosphere Bacterial Community Assembly in Wheat. Frontiers in Microbiology 10.

Korbie DJ, Mattick JS. 2008. Touchdown PCR for increased specificity and sensitivity in PCR amplification. Nature Protocols 3: 1452–1456.

Kuźniar A, Włodarczyk K, Grządziel J, Woźniak M, Furtak K, Gałązka A, Dziadczyk E, Skórzyńska-Polit E, Wolińska A. 2020. New Insight into the Composition of Wheat Seed Microbiota. International Journal of Molecular Sciences 21: 4634.

Larran S, Perelló A, Simón MR, Moreno V. 2002. Isolation and analysis of endophytic microorganisms in wheat (Triticum aestivum L.) leaves. World Journal of Microbiology and Biotechnology 18: 683–686.

Larran S, Perelló A, Simón MR, Moreno V. 2007. The endophytic fungi from wheat (*Triticum aestivum* L.). World Journal of Microbiology and Biotechnology 23: 565–572.

Latz MAC, Kerrn MH, Sørensen H, Collinge DB, Jensen B, Brown JKM, Madsen AM, Jørgensen HJL. 2021. Succession of the fungal endophytic microbiome of wheat is dependent on tissue-specific interactions between host genotype and environment. Science of The Total Environment 759: 143804.

Mascot-Gómez E, Flores J, López-Lozano NE. 2021. The seed-associated microbiome of four cactus species from Southern Chihuahuan Desert. Journal of Arid Environments 190: 104531.

Matanguihan JB, Murphy KM, Jones SS. 2011. Control of Common Bunt in Organic Wheat. Plant Disease 95: 92–103.

Mavrodi DV, Mavrodi OV, Elbourne LDH, Tetu S, Bonsall RF, Parejko J, Yang M, Paulsen IT, Weller DM, Thomashow LS. 2018. Long-Term Irrigation Affects the Dynamics and Activity of the Wheat Rhizosphere Microbiome. Frontiers in Plant Science 9.

Mergoum M, Johnson J, Buck J, Buntin GD, Sutton S, Lopez B, Mailhot D, Chen Z, Bland D, Harrison S, et al.2022. A new soft red winter wheat cultivar ‘GA 08535-15LE29’ adapted to Georgia and the U.S. southeast region. Journal of Plant Registrations 16: 597–605.

Mitter B, Pfaffenbichler N, Flavell R, Compant S, Antonielli L, Petric A, Berninger T, Naveed M, Sheibani-Tezerji R, von Maltzahn G, et al.2017. A New Approach to Modify Plant Microbiomes and Traits by Introducing Beneficial Bacteria at Flowering into Progeny Seeds. Frontiers in Microbiology 8.

Munkvold GP. 2009. Seed pathology progress in academia and industry. Annual Review of Phytopathology 47: 285–311.

Nelson EB. 2018. The seed microbiome: Origins, interactions, and impacts. Plant and Soil 422: 7–34.

Newcombe G, Harding A, Ridout M, Busby PE. 2018. A Hypothetical Bottleneck in the Plant Microbiome. Frontiers in Microbiology 9.

Nicolaisen M, Justesen AF, Knorr K, Wang J, Pinnschmidt HO. 2014. Fungal communities in wheat grain show significant co-existence patterns among species. Fungal Ecology 11: 145–153.

Noel ZA, Roze LV, Breunig M, Trail F. 2022. Endophytic Fungi as a Promising Biocontrol Agent to Protect Wheat from *Fusarium graminearum* Head Blight. Plant Disease 106: 595–602.

Oksanen J, Simpson GL, Blanchet FG, Kindt R, Legendre P, Minchin PR, O’Hara RB, Solymos P, Stevens MHH, Szoecs E, et al.2022. vegan: Community Ecology Package.

Pirttilä AM, S S. 2011. Prospects and Applications for Plant-Associated Microbes. A Laboratory Manual, Part B: Fungi.

Pochon S, Terrasson E, Guillemette T, Iacomi-Vasilescu B, Georgeault S, Juchaux M, Berruyer R, Debeaujon I, Simoneau P, Campion C. 2012. The *Arabidopsis thaliana*-*Alternaria brassicicola* pathosystem: A model interaction for investigating seed transmission of necrotrophic fungi. Plant Methods 8: 16.

R Core Team. 2018. R: A language and environment for statistical computing. R Foundation for Statistical Computing, Vienna, Austria.

Ridout ME, Schroeder KL, Hunter SS, Styer J, Newcombe G. 2019. Priority effects of wheat seed endophytes on a rhizosphere symbiosis. Symbiosis 78: 19–31.

Robinson RJ, Fraaije BA, Clark IM, Jackson RW, Hirsch PR, Mauchline TH. 2016. Wheat seed embryo excision enables the creation of axenic seedlings and Koch’s postulates testing of putative bacterial endophytes. Scientific Reports 6: 1–9.

Rojas EC, Sapkota R, Jensen B, Jørgensen HJL, Henriksson T, Jørgensen LN, Nicolaisen M, Collinge DB. 2020. Fusarium Head Blight Modifies Fungal Endophytic Communities During Infection of Wheat Spikes. Microbial Ecology 79: 397–408.

Russel J. 2022. MicEco: Various functions for microbial community data.

Schardl CL, Leuchtmann A, Spiering MJ. 2004. Symbioses of grasses with seedborne fungal endophytes. Annual Review of Plant Biology 55: 315–340.

Schlatter DC, Yin ChunTao, Hulbert S, Paulitz TC. 2020. Core rhizosphere microbiomes of dryland wheat are influenced by location and land use history. Applied and Environmental Microbiology 86: e02135–19.

Schulz B, Boyle C. 2005. The endophytic continuum. Mycological Research 109: 661–686.

Scibetta S, Schena L, Abdelfattah A, Pangallo S, Cacciola SO. 2018. Selection and Experimental Evaluation of Universal Primers to Study the Fungal Microbiome of Higher Plants. Phytobiomes Journal 2: 225–236.

Shade A, Jacques M-A, Barret M. 2017. Ecological patterns of seed microbiome diversity, transmission, and assembly. Current Opinion in Microbiology 37: 15–22.

Shiferaw B, Smale M, Braun H-J, Duveiller E, Reynolds M, Muricho G. 2013. Crops that feed the world 10. Past successes and future challenges to the role played by wheat in global food security. Food Security 5: 291–317.

Sieber T, Riesen TK, Müller E, Fried PM. 1988. Endophytic Fungi in Four Winter Wheat Cultivars (Triticum aestivum L.) Differing in Resistance Against Stagonospora nodorum (Berk.) Cast. & Germ. =Septoria nodorum (Berk.) Berk. Journal of Phytopathology 122: 289–306.

Simonin M, Dasilva C, Terzi V, Ngonkeu ELM, Diouf D, Kane A, Béna G, Moulin L. 2020. Influence of plant genotype and soil on the wheat rhizosphere microbiome: evidence for a core microbiome across eight African and European soils. FEMS Microbiology Ecology 96: fiaa067.

Singh D, Mathur SB. 2004. Histopathology of seed-borne infections. Boca Raton, USA: CRC Press Inc.

Singh T, Sinclair, J.B. 1985. Histopathology of *Cercospora sojina* in Soybean Seeds. Phytopathology 75: 185.

Smith SD. 2019. phylosmith: an R-package for reproducible and efficient microbiome analysis with phyloseq-objects. Journal of Open Source Software 4: 1442.

Solanki MK, Abdelfattah A, Britzi M, Zakin V, Wisniewski M, Droby S, Sionov E. 2019. Shifts in the Composition of the Microbiota of Stored Wheat Grains in Response to Fumigation. Frontiers in Microbiology 10: 1098.

Somma S, Amatulli MT, Masiello M, Moretti A, Logrieco AF. 2019. Alternaria species associated to wheat black point identified through a multilocus sequence approach. International Journal of Food Microbiology 293: 34–43.

Sowell A, Swearingen B. 2021. *Wheat Outlook: December* 2021. USDA-ERS.

USDA-NASS. 2021. Small Grains 2021 Summary. United States Department of Agriculture National Agricultural Statistics Service.

Vannette RL. 2020. The Floral Microbiome: Plant, Pollinator, and Microbial Perspectives. Annual Review of Ecology, Evolution, and Systematics 51: 363–386.

Vujanovic V, Islam MN, Daida P. 2019. Transgenerational role of seed mycobiome – an endosymbiotic fungal composition as a prerequisite to stress resilience and adaptive phenotypes in Triticum. Scientific Reports 9: 18483.

Vujanovic V, Mavragani D, Hamel C. 2012. Fungal communities associated with durum wheat production system: A characterization by growth stage, plant organ and preceding crop. Crop Protection 37: 26–34.

Walsh CM, Becker-Uncapher I, Carlson M, Fierer N. 2021. Variable influences of soil and seed-associated bacterial communities on the assembly of seedling microbiomes. The ISME Journal 15: 2748–2762.

Wassermann B, Rybakova D, Adam E, Zachow C, Bernhard M, Müller M, Mancinelli R, Berg G. 2021. Studying Seed Microbiomes. In: Carvalhais LC, Dennis PG, eds. Methods in Molecular Biology. The Plant Microbiome: Methods and Protocols. New York, NY: Springer US, 1–21.

Zhao C, Liu B, Piao S, Wang X, Lobell DB, Huang Y, Huang M, Yao Y, Bassu S, Ciais P, et al.2017. Temperature increase reduces global yields of major crops in four independent estimates. Proceedings of the National Academy of Sciences 114: 9326–9331.

